# Development and optimisation of Influenza C and Influenza D pseudotyped viruses

**DOI:** 10.1101/2025.05.26.656172

**Authors:** Maria Giovanna Marotta, Martin Mayora Neto, Janet Daly, Meshach Maina, Pauline van Diemen, Helen Everett, Maria Stella Lucente, Michele Camero, Emanuele Montomoli, Claudia Maria Trombetta, Kelly da Costa, Nigel J. Temperton

## Abstract

To facilitate the study of influenza C (ICV) and influenza D (IDV) viruses, we generated lentiviral pseudotyped viruses (PVs) expressing the hemagglutinin-esterase fusion (HEF) glycoprotein from ICV (C/Minnesota/33/2015) and IDV (D/Swine/Italy/199724-3/2015, D/Bovine/France/5920/2014, and D/Bovine/Ibaraki/7768/2016). The production of these PVs was optimised using different amount of human airway trypsin-like (HAT) protease to enhance HEF maturation, and the transduction efficiency was evaluated in multiple cell lines. Using these PVs, we established a pseudovirus-based microneutralisation (pMN) assay to measure neutralising antibody responses and adapted an esterase activity assay to evaluate PV. Specific antisera neutralised PVs but failed to inhibit esterase activity. These findings confirm that ICV and IDV PVs provide a scalable, sensitive, and safe tool for antiviral screening, and sero-epidemiological research.

## Introduction

Influenza is an infectious respiratory disease caused by the influenza virus, an enveloped, negative-sense single-stranded RNA virus, belonging to the *Orthomyxoviridae* family. This family comprises 4 genera: *Alphainfluenzavirus, Betainfluenzavirus, Gammainfluenzavirus*, and *Deltainfluenzavirus*, each corresponding to a single species: influenza A virus (IAV), influenza B virus (IBV), influenza C virus (ICV), and influenza D virus (IDV), respectively (1). All influenza viruses have a segmented genome. The genome of IAV and IBV consists of eight segments encoding multiple proteins, including the viral RNA polymerase subunits, the surface glycoproteins haemagglutinin (HA), which facilitates viral entry, and neuraminidase (NA), involved in viral release. They also encode viral nucleoprotein (NP), matrix proteins (M1 and M2), non-structural protein 1 (NS1), and nuclear export protein (NEP) (2). In contrast, ICV and IDV viruses possess only seven genomic segments, reflecting the presence of a single envelope glycoprotein, the hemagglutinin-esterase fusion (HEF) protein. HEF combines the functions of both HA and NA, being responsible for receptor binding, receptor destruction, and membrane fusion (3). The HEF glycoprotein is synthesized as a precursor (HEF0). Upon recognizing target cells, the viral HEF protein is cleaved by proteases into HEF1 and HEF2 subunits. This cleavage is essential for infection, determining both viral pathogenicity and tissue tropism.

Among the four influenza species, IAV has the broadest host range, infecting birds, humans, pigs, cats, horses, and rodents. It is also responsible for the most severe diseases and is associated with significant morbidity and mortality (1). IBV primarily infects humans, and it can cause seasonal epidemics ranging from mild to severe illness (4). Less is known about ICV, first isolated in 1947 from a human infection (5). While most individuals develop antibodies against ICV early in life, indicating prior exposure, clinical disease caused by ICV is rare, with the highest detection rates observed in children with lower respiratory tract infections (6). Studies suggest that between 78% and 100% of individuals develop antibodies against ICV (7). Knowledge of the epidemiology and virology of ICV are limited, as the virus is difficult to grow in culture and commercial validated test kits are unavailable. Although humans are the primary reservoir, ICV has been shown to infect pigs (8) and cattle (9), with serological evidence also suggesting infection in dogs (10, 11). Based on the HEF gene sequence, ICV is classified into six lineages: C/Taylor (1967), C/Aichi (1991), C/Yamagata (2004), C/Mississippi (2004), C/Sao Paulo (including the S1 and S2 sublineages), and C/Kanagawa, which are currently in circulation (12, 13). In 2011, a virus resembling ICV was isolated from pigs in the United States (14) and later from cattle (15), which were found to be the primary reservoir. Phylogenetic analysis revealed that this strain shared only 53% overall amino acid identity with human ICV, leading to its classification as a new genus within the *Orthomyxoviridae* family, named IDV. Serological studies have also revealed IDV positivity in several animal species, such as horses, sheep, and other ruminants (16, 17, 18). Although IDV has not been associated with human illness, seropositivity has been observed (19), particularly in individuals with occupational exposure (20, 21). IDV has evolved into five distinct lineages: the two oldest, D/Oklahoma and D/660, are prevalent in the Americas and Europe, while the more recent D/Yama2016 and D/Yama2019 circulate in Japan (22, 23, 24).

The fifth lineage (D/ CA2019) was reported in California, USA, after reassortment of the P3 gene of D/OK lineage and the genome segments of D/660 lineage (25). Pseudotyped viruses (PVs) are replication-defective viral particles comprising a structural core from one virus and a lipid envelope containing the surface glycoproteins of the virus of interest, along with a reporter component for quantification (26, 27). Given the challenges of culturing ICV and understanding the zoonotic potential of IDV, the production of optimised ICV and IDV PVs offers significant advantages for studying these viruses. PVs are cost-effective and suitable for large-scale neutralisation assays, without the need for high biosafety-level laboratories. PVs are widely used to study various aspects of viral biology, including sero-epidemiology, vaccine efficacy, and the screening of antivirals or monoclonal antibodies (26). The primary aim of this study was to produce and optimise PVs for ICV, specifically the C/Minnesota/33/2015 strain (São Paulo lineage), and for IDV strains, including D/Swine/Italy/199724-3/2015 and D/Bovine/France/5920/2014 (both Oklahoma lineage) and D/Bovine/Ibaraki/7768/2016 (Yama2016 lineage) (Figure 1).

**Figure 1.**
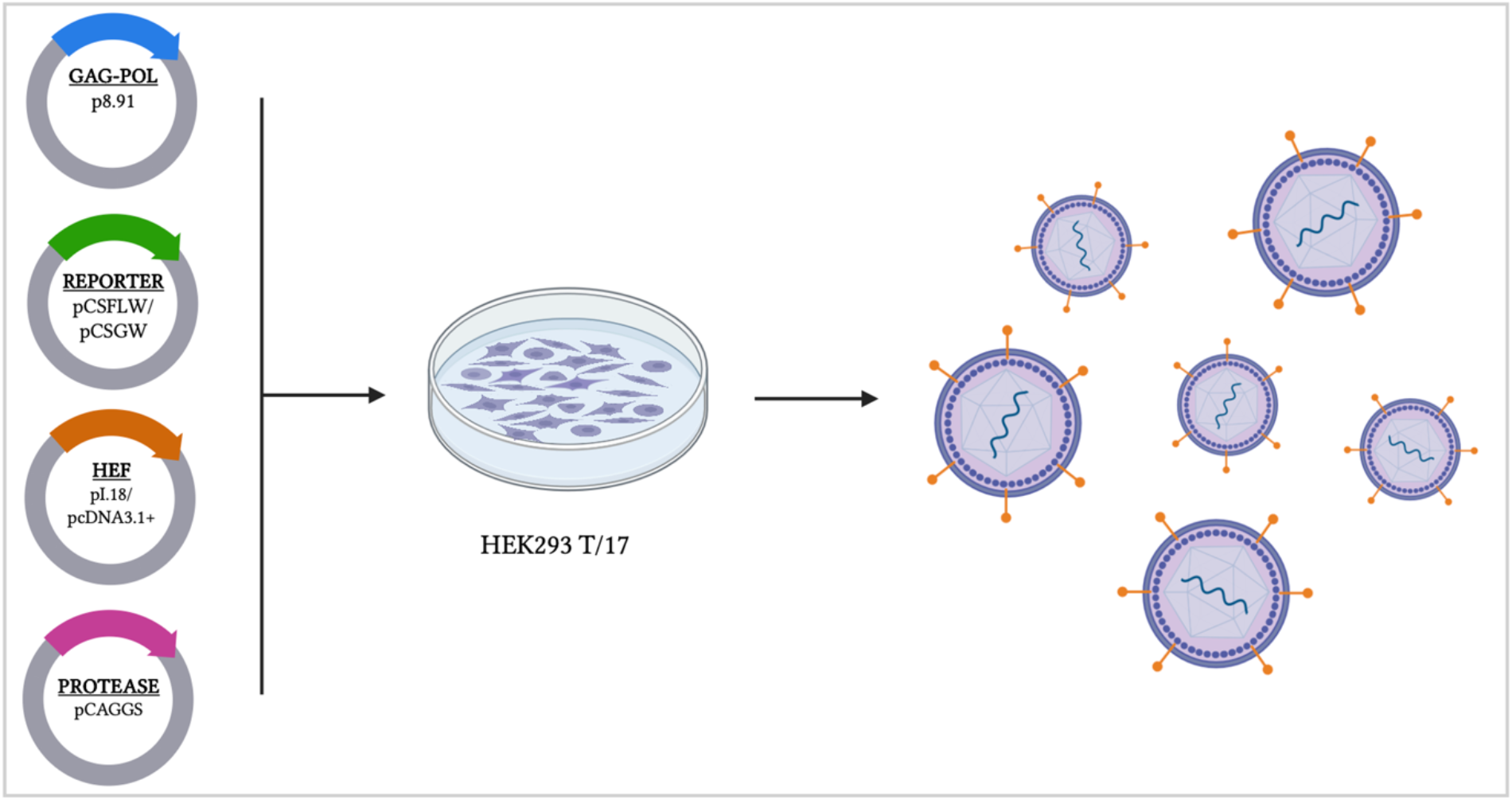
Graphical representation of ICV and IDV PVs production. The production involves a four-plasmid transfection method in HEK293 T/17 cell line: a plasmid encoding the gag-pol gene from HIV lentivirus for the viral core, a reporter plasmid (firefly luciferase or GFP), the plasmid encoding the HEF glycoprotein, and a protease-encoding plasmid to facilitate viral growth and HEF maturation. (Created in https://BioRender.com)

Herein, we show the development and optimisation of the production process, define the optimal target cell for use with these PVs, and then how to scale up production. We have evaluated the viral specific entry of these PVs both qualitatively via green fluorescent protein (GFP) and quantitatively using a luciferase reporter. We then demonstrate that these PVs can be used to assess the neutralisation capacity of serum samples. Additionally, we have adapted an esterase activity assay with the generated PVs to examine the HEF protein’s esterase activity.

## Materials and methods

### Cell line cultures

The human embryonic kidney 293T/17 (HEK293 T/17, ATCC: CRL-11268a) cells were used for transfection and tested as potential target cells. Swine testicular (ST) cells, madin–darby canine kidney II (MDCKII) cells, human hepatoma derived (Huh-7) cells were also tested as target cells. All the cell lines were maintained in complete medium Dulbecco’s Modified Eagle Medium (DMEM) (PANBiotech, P04-04510), with high glucose and GlutaMAX supplemented with 10% (*v*/*v*) heat-inactivated fetal bovine serum (PANBiotech, P30-8500) and 1% (*v*/*v*) penicillin–streptomycin (PenStrep) (Sigma-Aldrich, P4333) at 37 ^°^C and 5% CO_2_.

### Plasmids production and transformation

The HEF gene of ICV strain C/Minnesota/33/2015 (GenBank accession number: AST08292.1), designated as HEF_Minnesota_, was synthesized and subcloned into the mammalian protein expression plasmid (pcDNA3.1+) by GenScript, UK.

The HEF gene of IDV strain D/Swine/Italy/199724-3/2015 (GenBank accession number: KT592533.1), designated as HEF_Italy_, was synthesized by Thermo Fisher Scientific, UK and subcloned into the mammalian protein expression plasmid (pI.18) in house. The HEF pI.18 plasmid of D/Bovine/France/5920/2014 (GenBank accession number: MG720235.1), HEF_France_, was obtained by the methods described in Maina (2024) (28). The HEF gene of IDV strains D/Bovine/Ibaraki/7768/2016 (GenBank accession number: LC128433.1), designated as HEF_Japan_, was synthesized by GenScript, UK, and subcloned into pI.18 in house. Plasmids were transformed in chemically induced competent *E.coli* DH5*α* cells (Invitrogen 18265-017) via the heat-shock method. Plasmid DNA was recovered from transformed bacterial cultures via the plasmid mini kit (Qiagen 12125, Manchester, UK). All DNA extracts were quantified using UV spectrophotometry (NanoDrop™—Thermo Scientific, Paisley, UK).

### Pseudotyped virus (PV) production

For production of ICV and IDV PVs, transfection was performed as previously described (Figure 1) (29, 30). Briefly, 4 × 10^5^ cell/well of HEK 293T/17 cells in complete DMEM were seeded in a 6-well plate 24 hours before transfection and incubated at 37°C, 5% CO_2_ overnight. The next day, media was replaced, and cells were transfected using Opti-MEM™ (Thermo Fisher Scientific 1985062, Paisley, UK) and FuGENE^®^ HD transfection reagent (Promega E2312 Madison, USA) with the following plasmids: HEF_Minnesota_/ HEF_Italy /_HEF_Japan_ encoding plasmids in pI.18, p8.91 HIV gag-pol (gag-pol expression plasmid), luciferase (pCSFLW) or GFP (pCSGW) reporter plasmids. Plasmid expressing human airway trypsin-like protease (HAT) in pCAGGS was also included to provide protease for HEF maturation. All plasmid DNAs were mixed in Opti-MEM and FuGENE^®^ HD added dropwise followed by incubation for 15 min. The plasmid DNA-OptiMEM™ mixture was then added to the cells with constant swirling. The amounts of plasmid DNA and reagents used for transfection in a single well of a 6-well plate are summarised below (Table 1). Forty-eight hours post-transfection, supernatants were collected, passed through a 0.45 *μ*m cellulose acetate filter, and stored at −80°C. The VSV G PV and ΔEnvelope PV (where only the plasmids containing gag*-*pol and reporter gene were transfected into cells) were used to create positive and negative control PVs, respectively.

**Table 1.** Amounts of transfection components to produce ICV and IDV PVs.

| Solutions/Plasmids | Amount per well |
| --- | --- |
| OptiMEM™ | 100 μL |
| GAG-POL | 250 ng |
| Reporter | 375 ng |
| HEF | 250 ng |
| Protease | 125-500 ng |
| FUGENE® HD | 3 μL per μg of total plasmid DNA |

### Pseudotyped virus (PV) titration

Viral supernatants with luciferase reporter were serially diluted two-fold from 1:2 in a white 96-well plate (Thermo Fisher Scientific 13610, UK) in duplicate. Fifty *μ*L of target cells at a concentration of 1.75 × 10^4^ cell/well were added. Cell only control wells were also included on each plate. Plates were then incubated at 37°C, 5% CO_2_ for 48 h. After 48 hours, media was removed and 25 *µ*L of Bright-Glo^®^ (Promega, Madison, USA) luciferase assay substrate, in a 1:1 mix with Phosphate Buffer Saline (PBS), was added to each well. Plates were then read using the GloMax^®^ Navigator (Promega, UK) using the Promega GloMax^®^ Luminescence Quick-Read protocol. Viral pseudotype titre was then determined in relative luminescence units/mL (RLU/mL) using Excel.

Titration of GFP expressing PVs were performed in 96 well UV Transparent Plate (Thermo Fisher Scientific 8404, Paisley, UK) and were analysed by fluorescence microscope (ZOE™ Fluorescent Cell Imager; BioRad). The viral PV titre was determined in relative fluorescence units (RFU) using Excel, following manual counting of fluorescent cells. The number of GFP-positive cells was normalised to the volume of the virus used for infection. The obtained value was then multiplied by the dilution factor applied during sample preparation to calculate the final PV titre.

### Reference antisera

Immunoglobulin-depleted serum (Veterinary Lab Agency, PA0631) was used as a negative control, while Polyclonal Anti-Influenza Virus C/Taylor/1233/1947, (antiserum, Rooster) - NR-3132 (BEI Resources, NIAID, NIH) was reconstituted with 500 *µ*l of sterile dH_2_O and was used as a positive control for ICV-PVs. A serum sample from a pig vaccinated against D/Swine/Oklahoma/1334/2011 provided by Dr Pauline van Diemen (APHA), was used as a positive control for IDV-PV. Additionally, two serum samples from household adult dogs and two serum samples from cows, collected for research purposes in Apulia region (Italy), were also included. These samples were previously screened in house using the haemagglutination inhibition (HI) assay, with dog sera tested also against ICV (31) and cow sera against IDV. All samples were heat-inactivated at 56ºC for 30 minutes before use.

#### Ethics approval

*In vivo* studies were conducted in accordance with UK Home Office regulations under the Animals (Scientific Procedures) Act 1986 (ASPA) following ethical approval.

### Pseudotyped virus-based microneutralisation (pMN) assay

The pseudotyped virus-based microneutralisation (pMN) assay was performed in a white 96-well plate format. Briefly, antisera/sera were diluted 1:100 in DMEM in row A. A serial two-fold dilution was performed from row A to row H. Then, 50 *µ*L of PV at 2 × 10^7^ RLU/mL were added to each well. The serum-PV mixture was incubated for 1 hour at 37°C. After incubation, 50 *µ*L of ST cells at 1.75 × 10^4^ cell/well were added. The plates were then incubated at 37°C for 48 hours, 5% CO_2_. After the incubation period, all media was removed and 25 *µ*L of Bright-Glo™ Luciferase Assay System (Promega, Southampton, UK), in a mix 1:1 with PBS, were added to each well. Following a 5-minute incubation, the plates were read using a GloMax® Navigator luminometer (Promega, Southampton, UK) with the Promega GloMax® Luminescence Quick-Read protocol.

### Esterase activity assay

The enzymatic activity of the HEF glycoproteins from both ICV and IDV PVs was assessed using p-nitrophenyl acetate (pNPA), (N8130, Sigma–Aldrich) as a substrate. The pNPA is a reagent that in the presence of an esterase enzyme produces p-nitrophenol directly in proportion to enzymatic activity. Viral supernatants (100 *μ*L) of C/Minnesota/33/2015 and D/Swine/Italy/199724-3/2015 PVs were serially diluted in duplicate two-fold across a 96-well plate using either PBS or Bovine Serum Albumin (BSA) as buffers. To each well, 50 *μ*L of pNPA (diluted 1:10, 1:100, and 1:1000 in dH_2_O) was added and negative controls (only buffer and pNPA) were included. Optical density at 405 nm (OD_405_) was determined after different time of incubations and using the Tecan Sunrise™ microplate reader (Tecan Life Sciences, Switzerland) with Magellan™ data analysis software. Results were expressed as OD. Readings were normalised with the negative control and the dilution that resulted in 90% OD_405_ was selected as the PV dilution input for inhibition assays. Antiserum positive for ID_Italy_ was tested to determine its potential inhibition of esterase activity, starting at a dilution of 1:100 and with varying incubation times with PVs before readout: one hour, two hours, and overnight (18-20 hours).

### Statistical analysis

All statistical analysis was performed with GraphPad Prism Version 10.3.1 (GraphPad Software). PV titres were estimated using Excel™ software. The pseudotype titres obtained at each point in a range of dilution points were expressed as RLU/mL, and the arithmetic mean was calculated. To evaluate cell infectivity, an unpaired two-tailed Student’s test analysis was performed to calculate statistical significance (*, **, ***, and **** = *p* < 0.05, significantly differences; ns = not significant). Titres from pMN assays were firstly normalised to cell only (100% neutralisation) and PV only (0% neutralisation) controls. IC_50_ values were calculated by a non-linear regression model (log [inhibitor]) vs. normalised response-variable slope) (27). Results from enzymatic activity were normalised using Excel™ software and visualised using GraphPad Prism Version 10.3.1 (GraphPad Software).

## Results

### Optimisation of influenza C and D pseudotyped virus production and evaluation of pseudotyped virus-based microneutralisation (pMN) assay

The ICV and IDV PVs production was carried out by transfecting HEK293 T/17 cells, using a standard production protocol (Table 1) (29), and the PVs were harvested 48 hours post-transfection. Transduction of IC_Minnesota_ and ID_Italy_ was then performed in various cell lines (ST, MDCK II, HEK293 T/17, and Huh-7) (Figure 2A) using luciferase read out. The titre of IC_Minnesota_ was highest when ST cells were used as the target, with minimal variability across replicates (*p* = 0.0147). For ID_Italy_, transduction titres in HEK293T/17 and ST cells were comparable; however, ST cells showed lower variability (Figure 2A). Therefore, ST cells were selected as target cells for other IDV strains (ID_France_ and ID_Japan_). Additionally, the significantly higher titres of ICV and IDV-PVs compared to delta envelope (ΔEnvelope) confirm that viral-specific entry is required for infection. Previous studies have demonstrated that influenza viruses with a monobasic cleavage site can be proteolytically activated in cell culture by the addition of trypsin (32, 33, 34). Based on the work of our collaborators (28), in which several proteases were evaluated for their ability to activate the HEF glycoprotein of the ICV and IDV PVs, HAT was selected in this study as it showed the highest capacity to promote HEF maturation. As illustrated here (Figure 2B), the addition of HAT protease improved the titre of ICV-PVs compared to the delta protease (ΔProtease). However, using 125 ng of the HAT encoding plasmid resulted in a statistically significant increase of IC_Minnesota_-PV titre compared to the ΔProtease control (*p* < 0.05). As the optimal replication temperature for ICV is 33°C, due to its primary site of infection being the upper respiratory tract (35, 36), we also performed transfection of HEK293 T/17 cells at 33°C. After harvesting, ICV-PVs were transduced into ST cells at both 33°C and 37°C. The results, presented in Figure 2C, show a significant improvement in viral titre when PVs were transduced and titrated at 33°C, with a 1 log_10_ increase compared to those processed at 37°C (*p* < 0.0001). Additionally, a 2 log_10_ increase was observed when ICV-PVs were transfected at 33°C and transduced at 37°C, compared to those both transfected and transduced at 37°C (*p* < 0.0001). These findings align with previous studies (35, 36), confirming that ICV exhibits enhanced replication efficiency at 33°C. Moreover, ICV-PVs generated in this study closely mimic the biological behaviour of the wild-type virus, supporting their use as a reliable model for studying ICV entry and replication dynamics. As shown in Figure 2C, both ID_Italy_ and ID_France_, belonging to the D/Oklahoma lineage, have exhibited the highest PV titre using 250 ng of HAT, with statistically significant differences from ΔProtease control. In contrast, ID_Japan_ (Yama2016 lineage) reached a high titre with 500 ng of HAT but the t-test analysis did not indicate a significant difference from the ΔProtease control, suggesting that ID_Japan_ titre could be further optimised.

**Figure 2.**
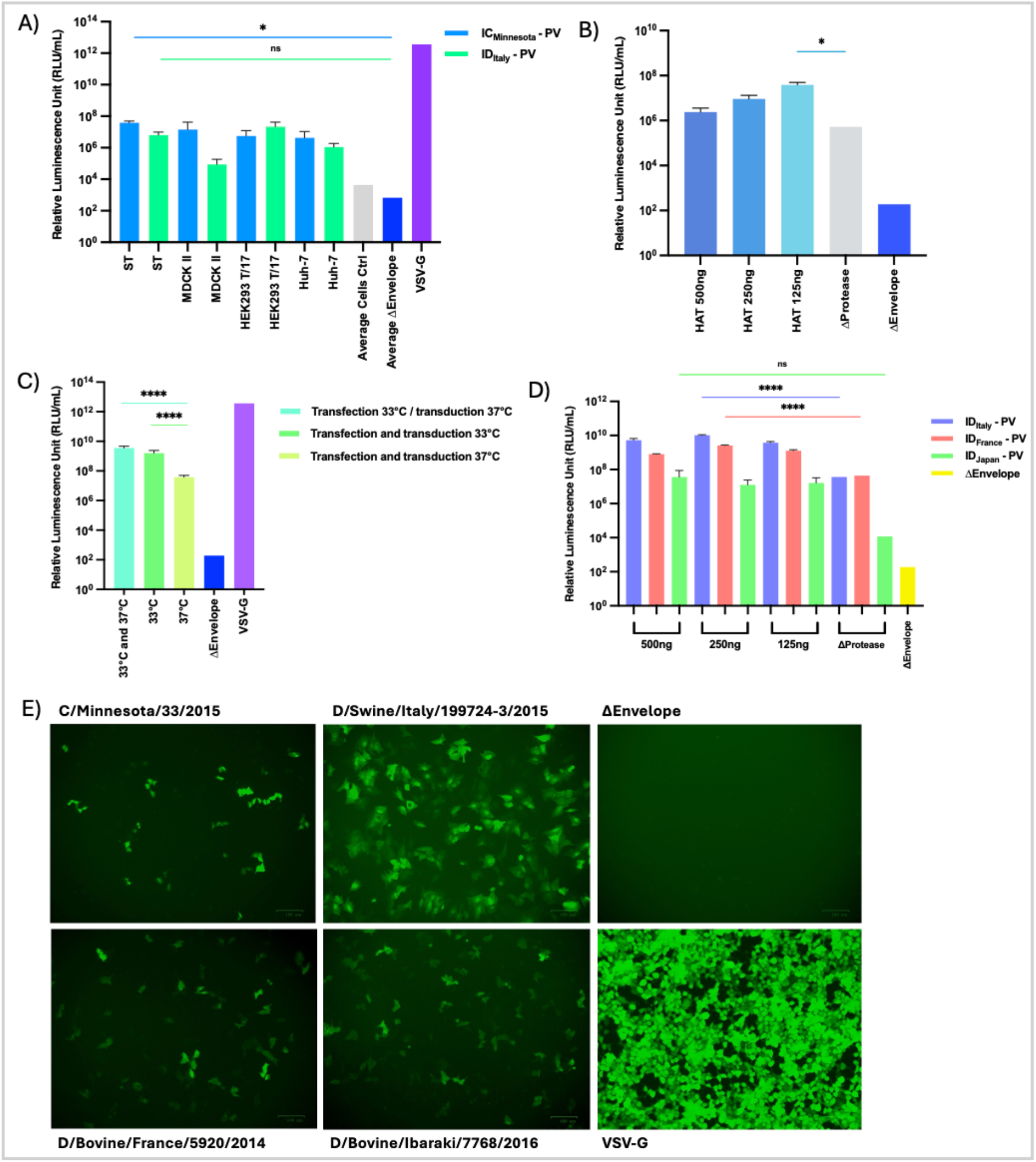
A) Cell lines susceptibility to ICV and IDV-PVs. Harvested, PVs were titrated using ST, MDCK II, HEK293 T/17, and Huh-7 cell lines for 48 hours to assess the optimal assay configuration. VSV-G PVs was used as a positive control and ΔEnvelope PVs (average from all cell lines) were included as negative control to confirm transduction from receptor specific entry. Statistical significance: *p* < 0.05 (*), ns = not significant. **B) Effect of HAT protease-encoding plasmid on IC_Minnesota_-PVs production**. IC_Minnesota_ -PVs were produced without protease-encoding plasmids (ΔProtease) or in the presence of varying amounts of HAT protease (500 ng, 250 ng & 125 ng per well) in a 6-well plate format. Harvested, PVs were then titrated on ST cells for 48 hours. ΔProtease and ΔEnvelope, PVs without glycoproteins, were used as negative controls. Statistical significance: *p* < 0.05 (*). **C) Effect of temperature on IC_Minnesota_-PV production.** HEK293 T/17 cells were transfected at either 37° or 33°C (5% CO_2_). PVs were harvested and transduced into ST cell for 48 hours at either 33°C or 37°C (5% CO_2_). Statistical significance: *p* < 0.0001 (****). **D) Effect of HAT protease-encoding plasmid on ID_Italy/France/Japan_-PVs production.** ID_Italy/France/Japan_-PVs were produced without protease-encoding plasmids (ΔProtease) or in presence of varying amounts of HAT protease (500 ng, 250 ng & 125 ng per well) in a 6-well plate format. Harvested, PVs were then titrated on ST cells for 48 hours. ΔProtease and ΔEnvelope, PVs without glycoproteins, were used as negative controls. Statistical analysis: *p <* 0.0001 (****), ns = not significant. Titres shown in the Figures 2A, 2B, 2C and 2D are mean RLU/mL values on a log-10 scale +/-standard deviation of two replicates. **E) Transfection efficiency of GFP-expressing PVs.** IC_Minnesota_-, ID_Italy_-, ID_France_-, and ID_Japan_-PVs were produced using a GFP reporter gene and transduced into ST cells. ΔEnvelope PVs (lacking glycoproteins) were used as negative controls, while VSV-G PVs (expressing VSV-G glycoproteins) served as positive controls. Images were captured 48 hours post infection using a fluorescence microscope. The scale at the right bottom represented as 100 *μ*m.

To confirm transduction efficiency and facilitate real-time visualisation, ICV and IDV PVs were additionally produced with a GFP reporter gene. Given their higher susceptibility to PV infection, ST cells were selected as the optimal target cell line, as described above. For IC_Minnesota_-PV 125 ng of HAT was used; for ID_Italy_ and ID_France_ -PV 250 ng of HAT has been employed, while 500 ng was used for ID_Japan_. Fluorescence images were captured 48 hours post-infection of ST cells using ZOE™ fluorescence microscope. As shown in Figure 2E, panels of IC_Minnesota_ and ID_Japan_ exhibited lower fluorescence signal compared to ID_Italy_ and ID_France_. These results are consistent with those obtained using the luciferase reporter gene (Supplementary, Figure 1S), confirming efficient production of these pseudotyped viruses.

To assess the inhibition susceptibility of IC_Minnesota_-PV in the pMN assay, we tested a specific reference antiserum C/Taylor/1233/1947 along with two sera from household adult dogs that had been previously tested in HI (31). Additionally, a positive antiserum from a pig vaccinated with D/Swine/Oklahoma/1334/2011 was used against ID_Italy_-PV, along with two cow sera previously tested in an HI test with the wild-type virus. A negative control serum was included for both ICV and IDV PVs. The pMN assay was configured following a protocol used for IAV and IBV experiments (37). The neutralisation capacity of the reference antiserum and dog sera against IC_Minnesota_-PV was compared to the negative control (pink color) (Figure 3A), showing a clear distinction between the two. Figure 3A illustrates the strong neutralisation capacity of the reference antiserum, and one dog serum (Sample No.98) that tested negative in HI (Supplementary Figure 2S) displayed low but detectable neutralising activity (IC_50_ = 78) (Figure 3B). This suggests that the pMN assay may be more sensitive than the HI assay for detecting neutralising antibodies. Similarly, Figure 3C shows that cow sera and antisera exhibited neutralising activity against ID_Italy_-PV. Cow sample No.184 shows stronger inhibition than Sample No.188 where no neutralisation capacity was observed (Figure 3D). These results align with those obtained in HI (Supplementary Figure 2S), reinforcing the ability of pMN to detect neutralising antibodies and highlighting its correlation with HI assay.

**Figure 3.**
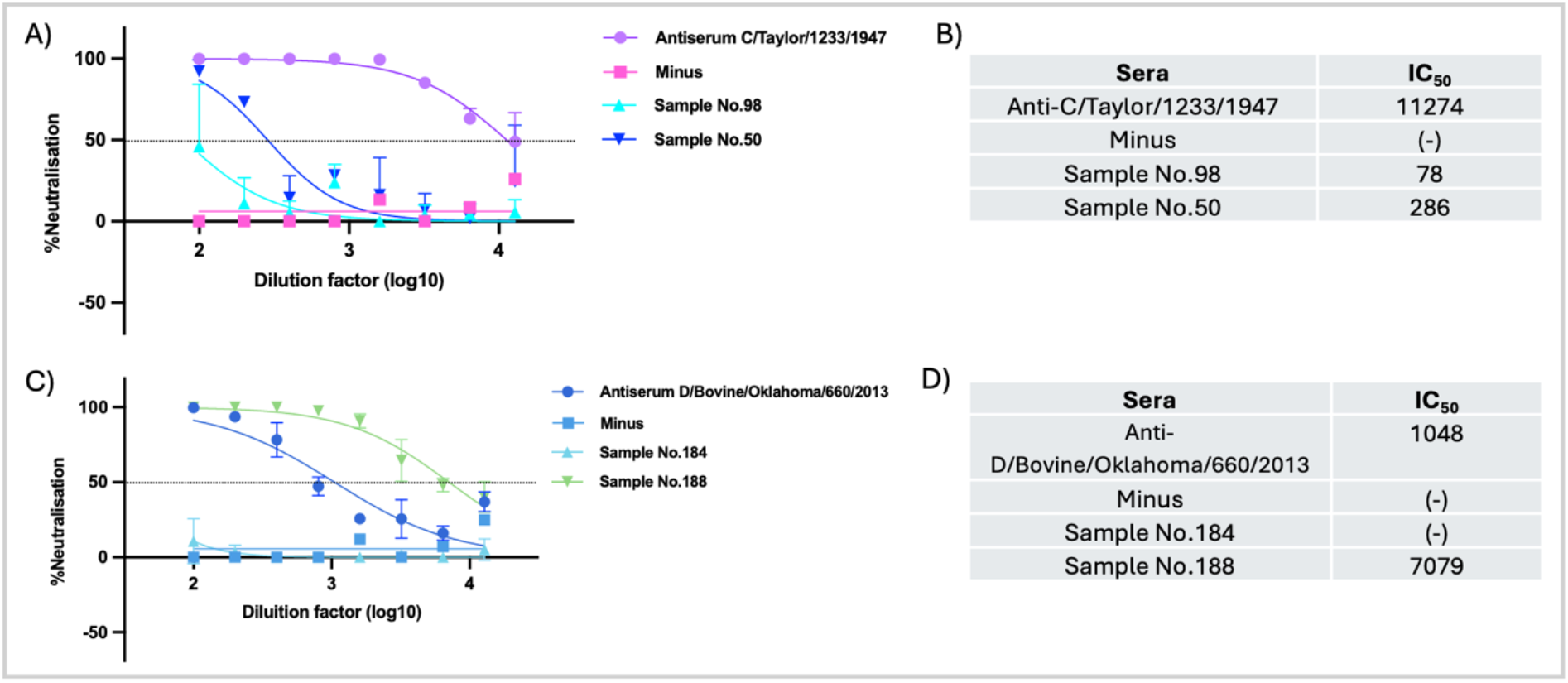
A) Neutralisation of IC_Minnesota_-PV using a luciferase-based pMN assay. Reference antiserum C/Taylor/1233/1947 (purple line), sera from two household adult dogs (light and dark blue lines), and a negative control (minus) (pink line) were tested. The reference antiserum was diluted 1:1000, while all sera were serially diluted two-fold from a starting dilution 1:100. A total of 1.0 × 106 RLU of PVs was then added to each well. The x-axis represents the sera dilution factor (log_10_), while the y-axis shows the percentage of neutralisation. Error bars indicate the standard deviation of two replicates per dilution. **B) IC_50_ values** from (A) using non-linear regression curve fit. **C) Neutralisation of ID_Italy_-PVs using a luciferase-based pMN assay.** The assay was performed similarly using reference antiserum D/Swine/Oklahoma/1334/2011 (dark blue line), two cow sera (light and dark green lines), and a negative control (minus) (light blue line). Sera were serially diluted two-fold from a starting dilution of 1:100, and total of 1.0 × 106 RLU of PVs was then added to each well. The x-axis represents the sera dilution factor (log_10_), while the y-axis shows the percentage of neutralisation. Error bars indicate the standard deviation of two replicates per dilution. **D) IC_50_ values** from (C) using non-linear regression curve fit.

### Optimisation of esterase activity assay to measure enzymatic activity of HEF glycoprotein

The HEF glycoprotein possesses esterase enzymatic activity, which facilitates the release of newly formed virion particles. The pNPA reagent has been demonstrated to effectively measure HEF enzymatic activity at different temperatures (38). To assess this activity in PVs, we adapted an esterase activity assay for IC_Minnesota_ and ID_Italy_-PVs. We tested PBS and BSA buffers as substrates to determine the most suitable buffer for the assay. Esterase activity was measured at 1 minute and subsequently every 5 minutes up to 30 minutes of incubation at 405 nm. Both ICV and IDV PVs exhibited detectable enzymatic activity (Figure 4). The esterase activity increased proportionally with incubation time in both PBS and BSA buffers. In contrast, negative controls using PBS or BSA alone did not show high activity.

**Figure 4.**
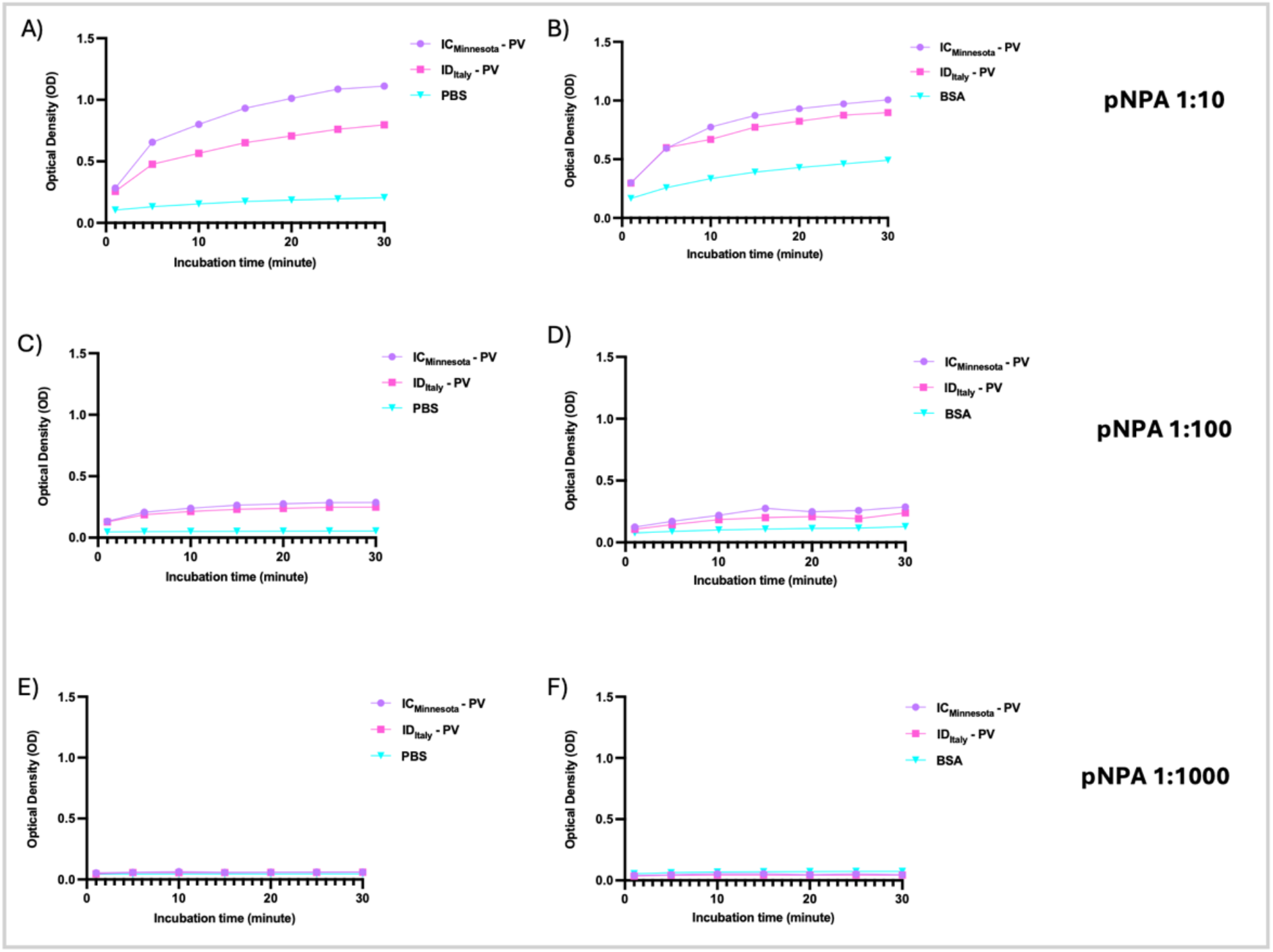
Esterase activity of HEF glycoproteins from IC_Minnesota_ and ID_Italy_ -PVs. Incubation time was measured from 1 min and at 5 minutes intervals for 30 minutes. P-nitrophenol acetate (pNPA) was used diluted 1:10 (A & B), 1:100 (C & D) and 1:1000 (E&F) in dH_2_O water. Only PBS or BSA were used as negative controls. The enzymatic activity is presented in Optical Density (OD) values measured at 405 nm.

The pNPA was initially tested at a starting concentration of 6.3 mg/mL, followed by serial dilutions (1:5, 1:10, 1:100, and 1:1000 in dH_2_O water) using both PBS (Figure 4A, C & E) and BSA (Figure 4B, D & F) buffers to determine the optimal concentration for enzymatic activity measurements. At undiluted and 1:5 dilution, pNPA did not yield reliable results. However, a consistent trend in enzymatic activity was observed across all other dilutions (1:10, 1:100, and 1:1000). The highest OD values were obtained using 1:10 dilution, indicating that this dilution is optimal for pNPA in this assay. Interestingly, a clear distinction between the IC_Minnesota_ and ID_Italy_ activity curves was observed when using PBS, whereas their curves were more closely aligned in BSA. This suggests that BSA enhances protein stability, minimising the differences between the two viruses in the assay (Figure 4A & B). Given that BSA is known to stabilise proteins by preventing non-specific binding through its blocking properties, it appears to be the most suitable buffer for this assay.

The esterase activity of HEF may serve as a potential drug target, like NA in IAV and IBV. For this reason, we further adapted the assay to evaluate the inhibition of HEF enzymatic activity by the antiserum D/Swine/Oklahoma/1334/2011. To determine the optimal PV dilution for inhibition assays, OD values were recorded at different incubation times, and the results were plotted to identify the dilution factor corresponding to 90% enzymatic activity (Figure 5A). Based on these findings, an incubation time of 20 minutes with pNPA before readout was selected for the inhibition assay in both ICV and IDV PVs. Next, ID_Italy_-PV OD values were plotted against dilution factors, and the curve was interpolated to determine the dilution factor yielding 90% esterase activity (Figure 5B). Based on these data, a dilution factor of 6 was chosen for further experiments. To assess potential HEF esterase inhibition, positive antiserum for ID_Italy_-PV was tested in BSA buffer. The starting dilution of the antiserum was 1:100, and incubation times were set at one hour, two hours, and overnight (18–20 hours) before adding pNPA. Each plate included pNPA and buffer (100% enzymatic activity control) and pNPA and PV (0% enzymatic activity control). Results were normalised, and the percentage of inhibition was calculated using a non-linear regression model (log [inhibitor] vs. normalised response-variable slope). Unfortunately, antiserum did not exhibit any inhibition of esterase activity, regardless of incubation time (Figure 5C). To evaluate the impact of incubation time on OD values, we analysed the control samples, ID_Italy_-PV and BSA. Figures 5D & E show an increase in OD values after 2 hours of incubation, while 1 hour of incubation resulted in greater data variability. However, when incubation was extended overnight (18–20 hours), OD values decreased (Figure 5F), suggesting that prolonged incubation may lead to enzymatic degradation or reduced assay sensitivity. Based on these results, an incubation time of 2 hours seems to be the optimal time to minimise measurement variation and maximise reliability. Therefore, 2 hours of incubation can be considered a standardized time point for future tests to ensure consistent OD measurements.

**Figure 5.**
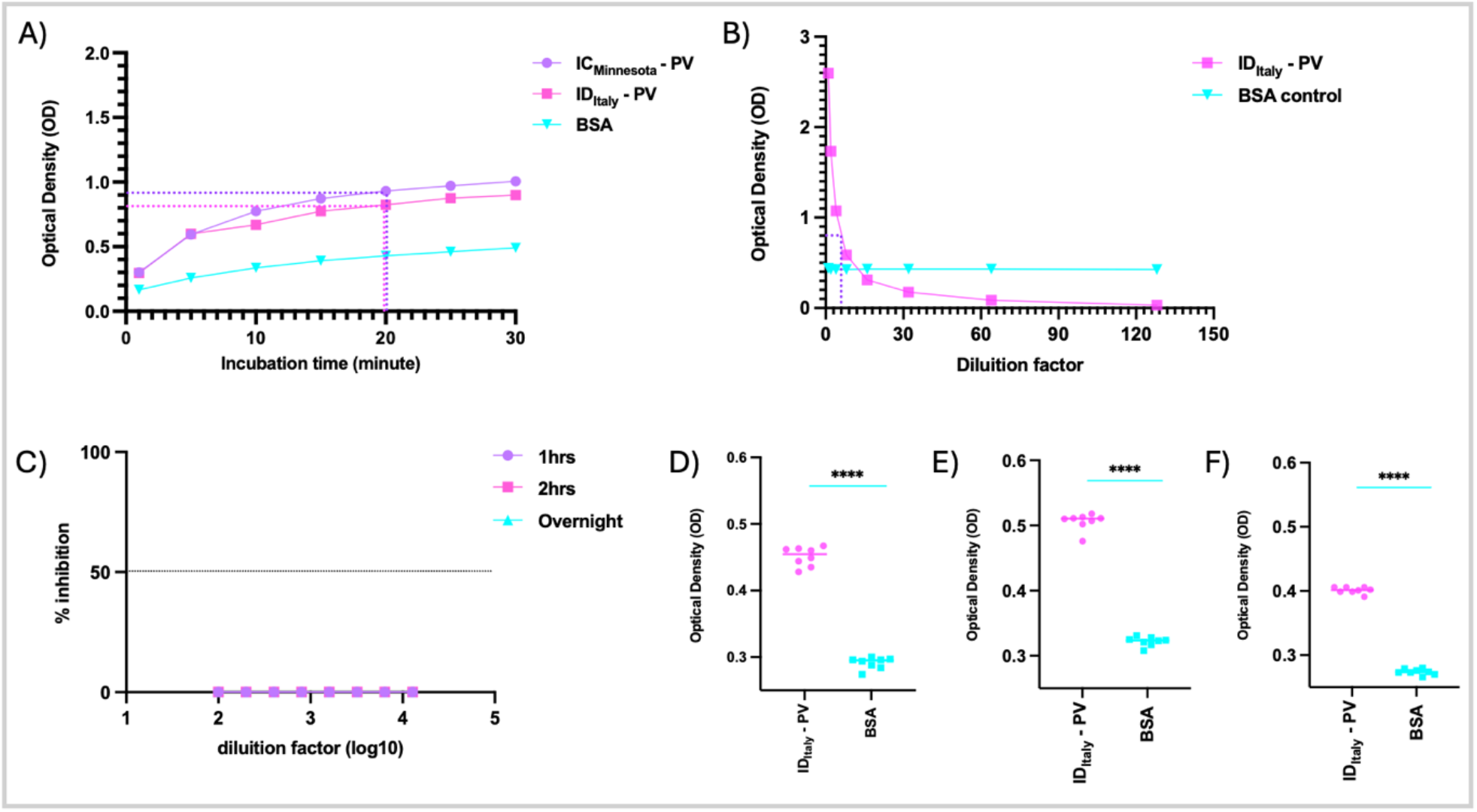
A) Determination of the optimal incubation time for esterase activity assay. The esterase activity of IC_Minnesota_-PV and ID_Italy_-PV was measured at different time points, starting from 1 minute and then every 5 minutes up to 30 minutes at 405 nm. OD values were recorded and plotted. BSA alone was used as a negative control. The curves were interpolated to determine the incubation time corresponding to 90% enzymatic activity. **B) Determination of the optimal PV dilution for inhibition assays.** OD values for ID_Italy_-PV were plotted against dilution factors, and the curve was interpolated to identify the dilution factor yielding 90% esterase activity. BSA was used as a negative control. **C) Evaluation of HEF esterase inhibition by ID_Italy_-PV antiserum.** An antiserum positive for ID_Italy_-PV was tested for potential inhibition of HEF enzymatic activity using BSA as a buffer. The starting dilution of the antiserum was 1:100, and incubation times were set at 1 hour, 2 hours, and overnight (18−20 hours) before adding pNPA. The inhibition percentage was calculated using a non-linear regression model. **D−F) Impact of incubation time on OD values in control samples.** Control samples (ID_Italy_-PV and BSA) were analysed after 1 hour (D), 2 hours (E), and overnight (F) of incubation. Statistical significance: *p* < 0.0001.

## Discussion

Although ICV and IDV are not currently classified as major threats to human health, it is essential to understand their biology and infection mechanisms to prevent and mitigate potential outbreaks. Despite their typically mild clinical presentations and limited pandemic potential—which have contributed to their underrepresentation in research—recent hospitalizations linked to ICV (39, 40) and increasing serological evidence of IDV exposure in humans (41) have raised concerns. The investigation of ICV and IDV rely on isolated virus, with PCR-based technique as the primary diagnostic method for ICV. Viral culture is particularly difficult for ICV due to its low *in vitro* replication efficiency, while for ICV and IDV the absence of a distinct cytopathic effect further limits their use in routine diagnostics and research. The pseudotyping system offers a safe and versatile approach to studying aspects of viral biology, providing a practical alternative method. PVs are non-replicative viral particles that undergo only a single round of infection (42), making them suitable for lower biosafety facilities and ideal for sero-epidemiological studies and antiviral screening. This approach has been widely adopted for various viruses, including influenza and SARS-CoV-2, as well as for high-risk pathogens such as Ebola and Lassa viruses, further demonstrating its utility in virology research (43, 44, 45, 46).

Our study demonstrates that PVs can effectively mimic ICV and IDV entry, supporting their use in the development of novel diagnostic tools and therapeutic strategies. A key advantage of this system is that HEF genes can be cloned into heterologous expression vectors (e.g., pI.18 and pcDNA3.1+), enabling expression once the viral sequence is known, without requiring virus isolation from clinical specimens. Moreover, the incorporation of a luciferase reporter gene allows for rapid, sensitive, and quantitative measurement of PV entry, highly adaptable to large-scale studies (47). An essential consideration in PVs production is the choice of target cells, as receptor distribution on the cell surface significantly influences viral entry efficiency. Our findings identified ST cells as the most susceptible to both ICV and IDV PVs, confirming their suitability for these applications. In line with the study of Maina (2024) (28) demonstrating the role of host proteases in cleaving HEF, we produced high titre ICV and IDV PVs using HAT protease. Notably, optimal protease concentrations varied between strains, suggesting that structural differences in HEF may impact protease recognition and cleavage efficiency, with potential implications for host adaptation, tissue tropism, and viral pathogenicity. Our findings also underscore the biological relevance of temperature in HEF maturation. Since ICV predominantly infects the upper respiratory tract (36), where the temperature is around 33°C, we tested this condition during PVs production. We observed a significant increase in ICV-PVs titre when transfections were carried out at 33°C. This reflects enhanced HEF folding and stability at lower temperatures, resulting in more PV particles. Interestingly, PVs transduction performed at 37°C yielded higher overall signal, likely due to increased metabolic activity and membrane fusion efficiency in target cells at physiological temperatures. We also employed GFP-transduced PVs, which enabled direct visualisation of infected cells, confirming that viral entry is mediated by specific interactions between the viral glycoprotein and host cell receptors. These findings were consistent with results obtained using luciferase-expressing PVs, demonstrating the reliability of both detection platforms. A major advantage of fluorescent protein PVs is their compatibility with microscopy-based screening and imaging platforms, allowing for automated quantification of infected cells without the need for additional substrates or specialized operator training.

PVs have proven to be an effective platform for measuring neutralising antibody responses across a range of host species, including mice, bats, sheep, and chickens (30, 48). In this study, we used ICV and IDV PVs to assess neutralising antibody responses in reference antisera, as well as in dog and cow serum samples previously tested by HI. All positive samples successfully neutralised the corresponding PVs, while negative sera showed no neutralising activity, confirming the assay’s ability to distinguish between positive and negative responses. Regarding dog and cow samples, the pMN assay demonstrated greater sensitivity than the HI assay. While HI is the gold standard for detecting antibodies targeting the globular head of HA, the pMN assay allows for the measurement of broadly neutralising antibodies directed against both the head and stalk regions of HA (38). This enhanced sensitivity may be due to the structural fidelity of the HEF glycoprotein expressed on the PV surface, which is presented in its native trimeric conformation. This allows for the detection of antibodies targeting conformational epitopes, including conserved regions not accessible in HI assay (49, 50, 51). Furthermore, pMN assays using PVs can capture the functional activity of a wider range of antibody isotypes, such as IgM, IgG, and IgA, unlike the HI assay, which primarily detects IgG (27, 42). These features highlight the broader immunological insights provided by pMN assays and support their use as an invaluable tool for evaluating antibody-mediated responses against ICV and IDV. Future work will focus on standardizing the ICV and IDV pMN assays by assessing performance parameters such as specificity, linearity, accuracy, precision, and robustness. These steps are essential to ensure assay reliability and support its implementation in the surveillance of zoonotic and emerging infections, as well as in the evaluation of universal influenza vaccines. In this context, and to contribute to universal influenza vaccine development and pandemic preparedness, we propose the inclusion of ICV and IDV PVs in a broader influenza pseudotype library. Given the challenges in isolating and characterising wild-type strains, especially for less studied viruses, PVs offer a flexible and powerful platform for serological analysis and pandemic risk assessment, particularly for ICV and IDV, which remain significantly less studied than IAV and IBV. PVs have also been successfully employed to evaluate antibody responses against NA of IAV and IBV (37). NA plays a critical role in the release of newly formed virions from infected cells (1, 2, 37, 47) and is the target of several licensed antiviral treatments. Similarly, the HEF glycoprotein of ICV and IDV not only mediates viral entry like HA but also exhibits esterase activity, which is essential for viral release (3, 4). This multifunctionality makes HEF an attractive target for antiviral strategies, as the inhibition of a single glycoprotein could potentially block both viral entry and release. To investigate this enzymatic function, we developed an esterase activity assay based on ICV and IDV PVs. To our knowledge, this is the first study to demonstrate the utility of a PV-based esterase assay for these viruses. We confirmed that both ICV and IDV PVs exhibit detectable enzymatic activity, further supporting their biological relevance. Although a standardized positive control is currently lacking, the assay is functional and suitable for measuring the enzymatic activity of various proteins, including the hemagglutinin esterase (HE) of coronaviruses (52, 53). This assay offers a practical and economic platform for assessing HEF function *in vitro* and holds potential for evaluating immune responses targeting the esterase domain, including neutralising antibodies that interfere with receptor-destroying activity. Moreover, it can be implemented in laboratories with lower biosafety requirements, providing an alternative to assays relying on wild-type virus. The system could also serve for screening HEF-targeting compounds, supporting the development of innovative therapeutic and prophylactic strategies against ICV and IDV.

In summary, due to the evolving influenza virus landscape and the ongoing risk of emerging strains, the development of novel and flexible assay systems is becoming increasingly critical. Traditional gold standard methods such as the HI assay, while still valuable, may not sufficiently capture the full range of viral functions or immune responses. To address this need, we demonstrate for the first time the utility of PV-based assay as a convenient and effective platform for investigating key aspects of ICV and IDV biology. Our study has defined the key components required for efficient ICV and IDV PVs production and optimised conditions to achieve high titres and assay performance. These conditions can be readily adapted to incorporate HEF from future ICV/IDV strains. Notably, we adapted the pMN system for use with ICV and IDV and developed a PV-based esterase activity assay to assess HEF function. Expanding the available PV-based platforms, such as the esterase activity assay described here, can enhance preparedness for future influenza threats and support the rapid evaluation of vaccines and antivirals in a safer and more adaptable context.

## Acknowledgements

We thank Juggragarn Jengarn for his contribution to the experimental work, part of which was previously published in Access Microbiology (2020) (https://doi.org/10.1099/acmi.ac2020.po0582).

## Author Contributions

**Conceptualization:** MGM, EM, CMT, KDC, NJT; **Formal Analysis and data curation:** MGM, KDC; **Investigation:** MGM, MMN, KDC; **Resources:** NJT, MC; **Supervision:** EM, CMT, KDC, NJT; **Visualization:** MGM; **Writing – original draft preparation**: MGM; **Writing – review and editing:** MMN, JD, MMM, PVD, HE, MSL, MC, EM, CMT, KDC, NJT.

