## Supplementary data for "Development and optimisation of Influenza C and Influenza D pseudotyped viruses"

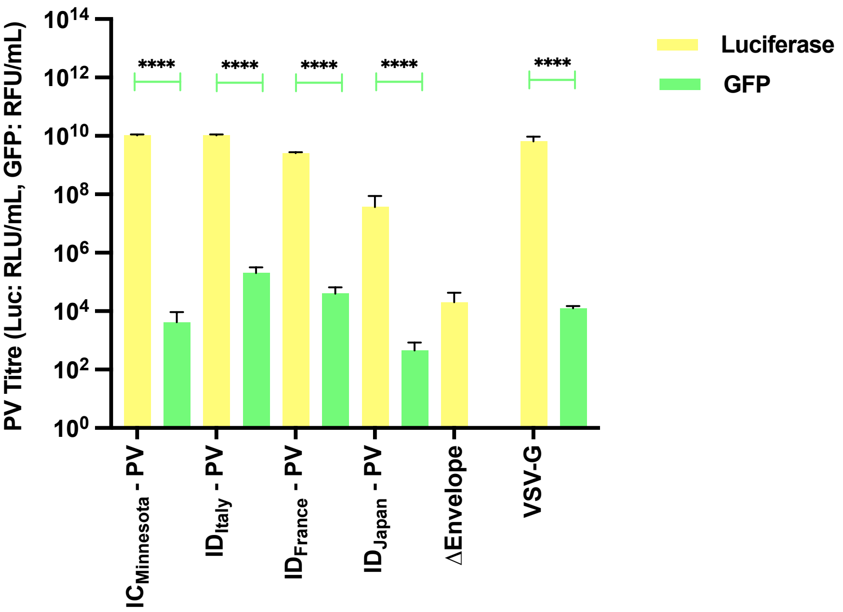


**Figure 1S. Comparison of PVs titration using Luciferase and GFP reporter genes.** PVs were produced with either luciferase or GFP as reporter genes, and titration was performed in ST cells to compare transduction efficiency. The bar graph represents the relative titres obtained from the two detection methods. Yellow bars indicate luciferase-based titration, while green bars represent GFP-based titration. The Y-axis represents the relative viral titre, measured in relative luminescence units per milliliter (RLU/mL) for luciferase-based titration and relative fluorescence units per milliliter (RFU/mL) for GFP-based titration**.** The results demonstrate differences in sensitivity and quantification accuracy between the two reporter systems, with luciferase generally yielding higher titres. These findings validate the use of both detection methods for assessing PV production efficiency and confirm consistency in viral entry measurements.

A)

| Dog sera | HI titre | $IC_{50}$ |
|---|---|---|
| Sample No.98 | 5 | 78 |
| Sample No.50 | 20 | 286 |

B)

| Cow sera | HI titre | $IC_{50}$ |
|---|---|---|
| Sample No.184 | 5 | (-) |
| Sample No.188 | 160 | 7249 |

**Figure 2S. A) Comparison of HI and pMN titres in dog sera.** Sample No.98 had a negative HI titre but showed a detectable neutralisation activity in pMN, suggesting a potential higher sensitivity of the pMN assay compared to HI. Sample No.50, which had a slightly higher HI titre, exhibited a higher $IC_{50}$, aligning results in pMN assay also indicating that pMN may detect some neutralisation activity in samples with low HI titres. **B) Comparison of HI and pMN titres in cow sera.** Sample No.184 showed negative HI titre and no detectable neutralisation in the pMN assay has been observed. Sample No.188, with a higher HI titre exhibited a much stronger neutralisation response. These results confirm the correlation between high HI titres and strong neutralisation activity, reinforcing the reliability of pMN as a complementary assay for detecting neutralising antibodies.
